# Optimisation of femtosecond dispersion compensation at the sample using side-view two-photon fluorescence imaging

**DOI:** 10.64898/2026.11.27.769678

**Authors:** Jakub Czubek, Grzegorz Orlicki, Jacek Łubiński, Małgorzata Szczerska, Filip Janiak

## Abstract

Efficient two-photon absorption depends on compensating the dispersion of the whole excitation path. Building on established fluorescence-based adjustment, we describe a practical procedure that uses side-view images of fluorescence in a dye cuvette, independently of the microscope’s scanning and detection, with reference records, stability warnings and open analysis software. For a 1045 nm laser, maxima lay between 21 and 24 screw turns in 11 of 13 sweeps. For a 920 nm laser, fluorescence maxima at the sample differed by 1.63 kfs² on average from autocorrelation minima before the scan optics. The recordings also show limitations from signal instability and proximity to clipping. The procedure targets two-photon microscopes for in vivo imaging, such as of neuronal activity, where tissue heating limits the usable power; it may also apply to photostimulation and polymerisation paths.

## 1. Introduction

Two-photon absorption underlies two-photon microscopy, the method of choice for imaging deep in scattering tissue [Denk et al., 1990; Zipfel et al., 2003; Helmchen and Denk, 2005], and two-photon polymerisation, which writes three-dimensional structures with sub-diffraction features [Maruo et al., 1997; Kawata et al., 2001]. Much effort goes into larger fields of view [Sofroniew et al., 2016; Stirman et al., 2016; Janiak et al., 2022], faster imaging [Wu et al., 2020], higher resolution through adaptive optics [Ji et al., 2010] and functional micro-devices [Gissibl et al., 2016; Trautmann et al., 2019] (for a review, see [Hoover and Squier, 2013]). All of these rely on efficient two-photon absorption at the focus. Its rate scales with the square of the instantaneous intensity, so at fixed average power and spectrum it is inversely proportional to the pulse duration at the sample [Pang et al., 2009]. Common transmissive microscope optics add positive group-delay dispersion (GDD) in this wavelength range: high-numerical-aperture objectives contribute 2000–6500 fs² [Guild et al., 1997] and complete systems several thousand to tens of thousands of fs² [Field et al., 2010; Palczewska et al., 2018; Lees et al., 2025]. An under-compensated pulse has to be offset with higher average power, which in tissue is limited mainly by heating [Podgorski and Ranganathan, 2016; Schmidt and Oheim, 2020], as in two-photon imaging of the mouse brain and the larval zebrafish retina [Podgorski and Ranganathan, 2016; Zhou et al., 2020; Bartel et al., 2021].

The pulse can be characterised at or near the focus by interferometric autocorrelation with a two-photon detector [Guild et al., 1997; Müller et al., 1995; Brakenhoff et al., 1995], frequency-resolved optical gating at full aperture [Fittinghoff et al., 1998; Fittinghoff et al., 1999], multiphoton intrapulse interference phase scan [Lozovoy et al., 2004; Xi et al., 2009] or dispersion scan [Miranda et al., 2012]. These methods require an interferometer, a nonlinear crystal with a spectrometer, a pulse shaper or a modification of the microscope, and commercial autocorrelators placed before the microscope miss the scan optics and the objective [Kim and Vogel, 2020]. In two-photon polymerisation, the pulse duration at the focus affects polymerisation and damage thresholds and has been measured with autocorrelators adapted to high-NA objectives [Samsonas et al., 2023].

In practice, compensation can be adjusted by maximising a nonlinear signal, a principle established for adaptive pulse compression [Yelin et al., 1997] and used in microscopy to maximise two-photon fluorescence [Field et al., 2010; Müller et al., 1998]. A community protocol describes stepping GDD while recording the mean pixel value of a fluorescent reference through the microscope’s photomultiplier [Lees et al., 2025]. Side-view camera detection has also been used: Saidi and Shtrahman imaged fluorescence through a lateral cuvette window and adjusted each laser’s compensator to maximise the brightest image pixel before comparing two-photon excitation efficiencies [Saidi and Shtrahman, 2023]. Thus, side-view fluorescence-based compensation is an established approach that does not require the microscope’s scanning or fluorescence detection. A practical acquisition and analysis procedure can help users interpret compensator sweeps and recognise instability, clipping and changes in sample geometry. Such documentation complements broader efforts to improve quality assessment and reproducibility in microscopy [Corbin et al., 2014; Nelson et al., 2021; Boehm et al., 2021; Faklaris et al., 2022; Janiak et al., 2026].

Watching two-photon fluorescence from the side is one of the oldest pulse diagnostics [Giordmaine et al., 1967], and direct imaging of excitation volumes has been used to characterise two-photon scan geometries [Janiak et al., 2022]. Building on this established geometry and its use for compensation adjustment [Saidi and Shtrahman, 2023], we develop a practical sweep procedure using a dye cuvette, a parked beam and a side-view camera with a compact lens. The contribution is the acquisition and interpretation workflow: reference records during sweeps, practical stability warnings, comparison of observed maxima between sessions, and documentation of failure modes. We report repeated sweeps on a 1045 nm laser with a mechanical compensator, comparison with upstream autocorrelation on a 920 nm fibre laser with software-controlled compensation, and observations of heating, sample displacement, reflections and sample damage. A typical use is a routine check of a two-photon microscope for in vivo imaging, for example of neuronal activity in the mouse brain or the zebrafish retina [Podgorski and Ranganathan, 2016; Zhou et al., 2020; Bartel et al., 2021], after a change of objective, wavelength or beam path, where recovering signal without raising the average power limits tissue heating. Open analysis software with a synthetic verification dataset and step-by-step instructions (Supplement 1) accompany the paper.

## 3. Results

### 3.1. Measurement geometry

Focusing into fluorescein produced a localised fluorescence spot inside the cuvette, clearly separated from the walls and the liquid surface and visible to the side-view camera (Fig. 1B). Because the beam is parked, the signal and background ROIs stay fixed for the whole sweep, and only the compensator setting changes between records (Fig. 1A,C,D).

**Fig. 1.**
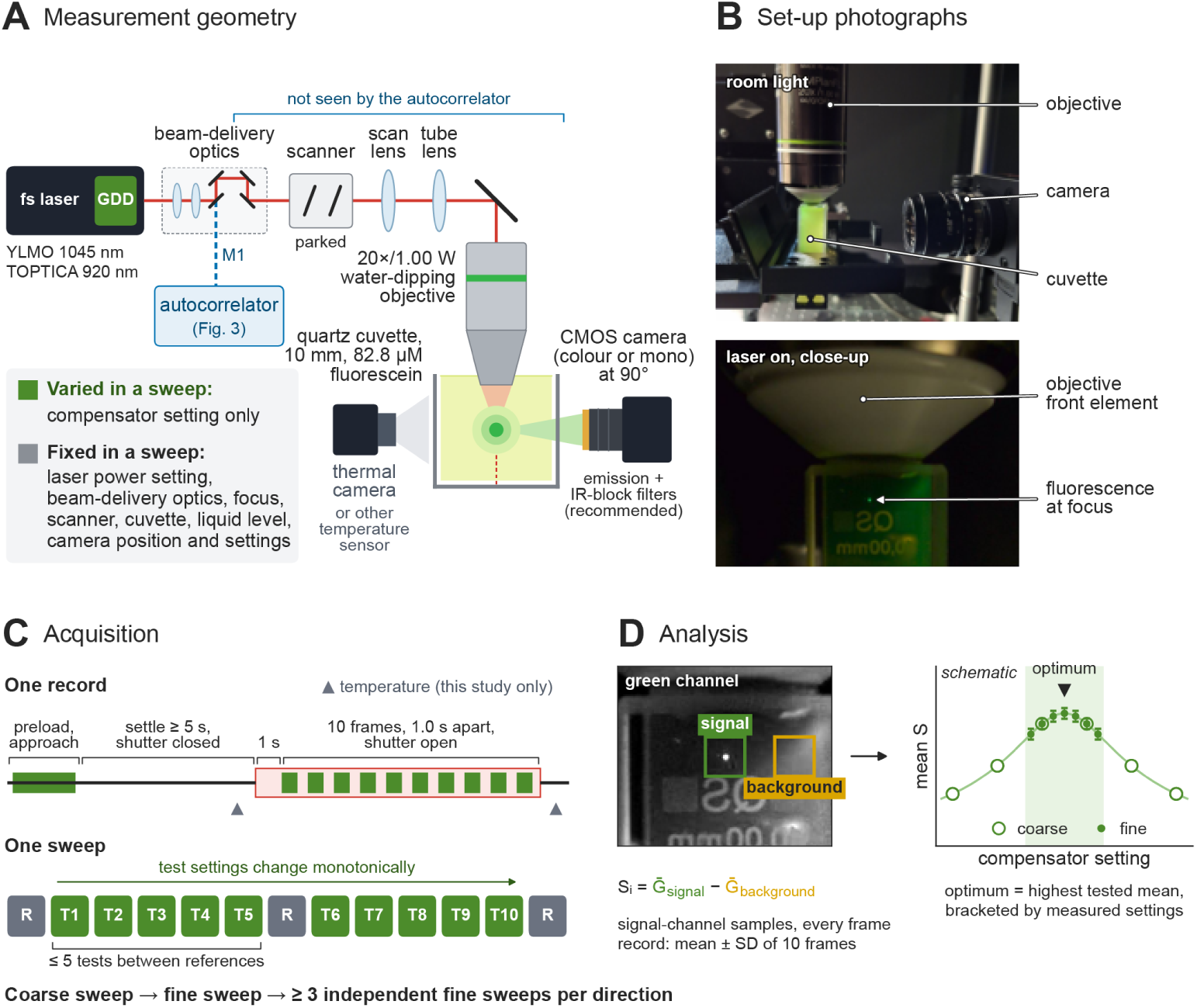
Measurement geometry, acquisition sequence and analysis for fluorescence-based dispersion optimisation. **(A)** Excitation path and detection geometry of the example configuration used in this study (schematic, not to scale). **Lasers.** Two femtosecond lasers were used: YLMO (1045 nm), with dispersion set by a mechanical compensation screw in turns, and TOPTICA (920 nm), with software-controlled GDD in fs². In Figs 3–5, the lasers are referred to by wavelength. **Beam path.** The beam passes through the beam-delivery optics (beam expander and steering mirrors), the parked scanner, the scan lens and the tube lens, and is focused by a 20×/1.00 W water-dipping objective (Olympus XLUMPlanFL) into 82.8 µM fluorescein (pH 9.0) in a 10 mm quartz cuvette (Hellma QS). The cuvette is filled to the rim, giving a flat liquid surface, and the objective is immersed in the solution without touching the walls of the cuvette. **Detection.** A colour CMOS camera (Thorlabs CS165CU) records the fluorescence at 90° to the excitation axis. No additional emission or IR-blocking filters were used in this study (the camera has a built-in IR-blocking filter); the protocol recommends them. In this study, a thermal camera also monitored sample temperature; another temperature sensor can be used, and the protocol does not require this measurement (see Fig. 4A,B). **Autocorrelator.** For the measurements in Fig. 3, the beam was re-routed at mirror M1, after the beam expander, to an autocorrelator (blue dashed path). The bracket marks the elements after M1, which the autocorrelator measurement does not include. **Fixed and varied.** Within a sweep, only the compensator setting (green) is varied; all other settings (grey) are kept fixed. **(B)** Photographs of the arrangement under room light (top) and during excitation (bottom). The arrow marks the localised fluorescence at the excitation focus inside the cuvette. **(C) Recommended acquisition sequence.** The panel labelled “Acquisition” illustrates the proposed protocol (Supplement 1), rather than the timing of the recordings analysed here; the acquisition used in this study is described in Section 2.4. **One record.** The compensator is moved to a preload position and then approaches the target setting from a fixed side to eliminate backlash. After at least 5 s of settling with the shutter closed, the shutter is opened. After a 1 s pre-exposure, ten frames are acquired at a 1.0 s frame-start interval. Grey triangles indicate optional temperature readings; temperature measurements are described in Section 2.8. **One sweep.** Reference records (R, grey) at a fixed compensator setting are acquired at the start, after no more than five test records (T, green), and at the end. Test settings change monotonically. A coarse sweep brackets the maximum, and at least three independent fine sweeps are then acquired per direction. **(D) Analysis. Left.** Green channel of the close-up image in (B), contrast-stretched for display, with fixed signal (green) and background (yellow) regions of interest; ROI positions are illustrative. For each frame i, the corrected signal is Sᵢ = Ḡ_signal_ − Ḡ_background_, where Ḡ is the mean of the signal-channel samples in the ROI: green Bayer sites for native colour-sensor data, channel 2 for linear RGB data, or all pixels for monochrome data. Each record is summarised as the mean ± SD of its ten frames. **Right.** Schematic sweep. Coarse settings (open circles) bracket the maximum, and fine settings (filled circles, mean ± SD) resolve it. The optimum is the highest tested mean of a sweep and must be bracketed by measured settings.

### 3.2. Consistency of the maximum with a mechanical compensator

Across 13 sweeps in nine sessions with the 1045 nm laser, the positions of the observed maxima were consistent between sessions (Fig. 2A). Excluding one sweep with a horizontal cuvette and one with sample displacement, the maxima lay between 21 and 24 turns, about 1.7 kfs² assuming a linear compensator. In the sweeps of 8 October, the maximum-pixel measure and the protocol metric showed similar sweep shapes, with the same broad maximum between positions 22 and 25 (r = 0.97 for both directions; Fig. 2B,C). The maximum-pixel measure compressed the contrast (0.89–1 compared with 0.73–1) because the brightest pixels approached the 8-bit ceiling.

**Fig. 2.**
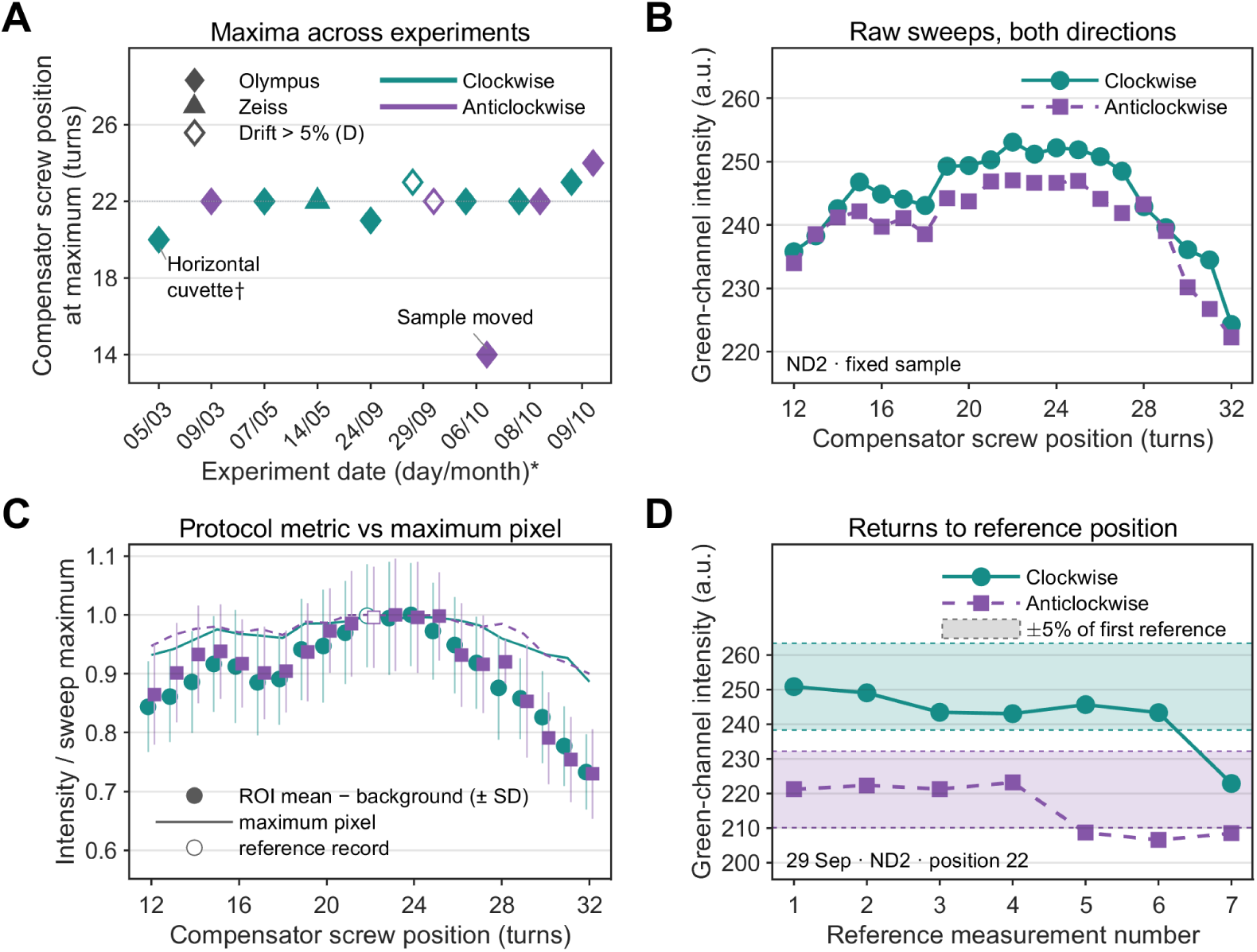
Reproducibility of fluorescence-based dispersion optimisation with the 1045 nm laser (YLMO). (A) Compensator screw positions at the maximum green-channel fluorescence in 13 sweeps across nine experimental sessions. The asterisk after the horizontal-axis label refers to dates being displayed as day/month, with sessions equally spaced rather than separated in proportion to elapsed time. Colours indicate clockwise (turquoise) and anticlockwise (purple) rotation; diamonds and triangles denote Olympus and Zeiss objectives, respectively. One turn of the screw corresponds to about 0.58 kfs² (30 000 fs² over 52 turns). Measurements with a horizontal cuvette or with sample displacement (Fig. 5A) are annotated; in the horizontal-cuvette measurement (†), the first of the series, one point (position 20, the maximum) reached the 8-bit ceiling (255) and may be saturated. Apart from these two measurements, peak positions ranged from 21 to 24 turns; the dotted line marks position 22. Open symbols mark the two sweeps of 29 September shown in (D), whose reference values changed by more than the ±5% warning level later adopted in the protocol. (B) Raw fluorescence intensity during clockwise and anticlockwise sweeps on 8 October with the ND2 filter and fixed sample geometry. (C) The same sweeps evaluated with the protocol metric (background-corrected ROI mean, Fig. 1D; points, mean ± SD of ten frames) and with the maximum-pixel measure used in (A) and (B) (thin lines), each normalised to its own maximum. Both measures show the same broad maximum between positions 22 and 25 (correlation between the two measures r = 0.97 for both directions). The maximum-pixel measure compresses the contrast (0.89–1 compared with 0.73–1) because the brightest pixels approach the 8-bit ceiling. Open symbols, reference record at position 22; points are offset horizontally by ±0.15 for clarity. The records exceed the proposed within-record drift warning level (reported within-record signal change over ten consecutive video frames, 16–22%; this subsecond measure is distinct from the minute-scale trends in Fig. 4) and do not meet the recommended headroom. They are shown to compare the two signal metrics; their agreement does not establish the accuracy of the inferred optimum. (D) Repeated returns to reference position 22 during the clockwise and anticlockwise sweeps of 29 September. Shaded bands with dashed limits mark ±5% of the first reference value of each sweep. The reference values changed by up to about 11% (clockwise) and 7% (anticlockwise), more than the ±5% warning level later adopted in the protocol, illustrating why the protocol includes repeated reference measurements. Seven reference measurements were acquired per sweep; measurement number denotes acquisition order rather than equally spaced time intervals. In B–D, circles and solid lines indicate clockwise sweeps, and squares and dashed lines anticlockwise sweeps; points are individual measurements and lines are guides to the eye, without fitting or smoothing. In A, B and D, intensity is the maximum green-channel pixel value of the ten-frame mean image (arbitrary units; 8-bit Motion-JPEG video), rather than the background-corrected ROI mean of Fig. 1D, and values close to the 8-bit ceiling (255) do not meet the 95% headroom recommendation of the protocol.

Repeated returns to the reference position revealed changes of up to 11% (clockwise) and 7% (anticlockwise) within a sweep (Fig. 2D), and the protocol-metric records of 8 October showed within-record signal changes reported as 16–22% across ten consecutive frames (about 0.3 s; the warning level refers to frames 1 s apart) and approached the 8-bit ceiling. These subsecond changes are not equivalent to the slow trends measured over minutes in Fig. 4, and their cause was not resolved. These recordings exceed the proposed stability warning levels and headroom recommendation. They illustrate the limitations encountered during method development and motivated the quality checks in Table 1. Although their maxima occur within a similar range of compensator settings, drift and proximity to clipping limit the precision with which a unique optimum can be assigned.

**Table 1.** Practical quality checks and recommended acquisition conditions.

| Criterion | Recommended condition or warning level | What it detects |
|---|---|---|
| Corrected signal | Positive mean | Non-positive background-corrected mean; a positive mean alone does not establish detection above noise |
| Temporal noise | Coefficient of variation of the ten frames ≤ 5% (author-selected warning level) | Unstable signal within a record |
| Within-record drift | \|Mean of the last three frames − mean of the first three\| ≤ 5% of the ten-frame mean (author-selected warning level) | Bleaching, heating or power drift during a record |
| Headroom | Maximum code in either ROI ≤ 95% of the verified clipping code | Clipping |
| Camera settings | Exposure, gain and gamma fixed and logged; automatic settings off | Uncontrolled changes in signal amplitude |
| Reference stability | Every reference within ±5% of the first (author-selected warning level) | Drift across the sweep (about 2.5% in power if the signal is quadratic) |
| Reference spacing | References at start and end, at most five tests between them | Undersampled drift |
| Bracketing | Highest mean not at a range boundary | Maximum outside the tested range |
| Repeats | At least three fine sweeps per direction are recommended to assess repeatability | Irreproducible optimum |

### 3.3. Comparison with autocorrelation

For the 920 nm laser, fluorescence sweeps at three power settings in separate sessions gave fitted maxima of −2.34, −1.91 and −2.40 kfs², a mean of −2.22 ± 0.26 kfs² (n = 3; Fig. 3A,B). Autocorrelation at mirror M1 gave pulse-width minima of −0.56 and −0.63 kfs², a mean of −0.59 ± 0.05 kfs² (n = 2; Fig. 3C). The maximum of the fluorescence signal at the sample therefore lay 1.63 kfs² from the minimum pulse duration measured before the scan optics (Fig. 3D,E); the sign is consistent with the additional positive dispersion of the elements after M1.

**Fig. 3.**
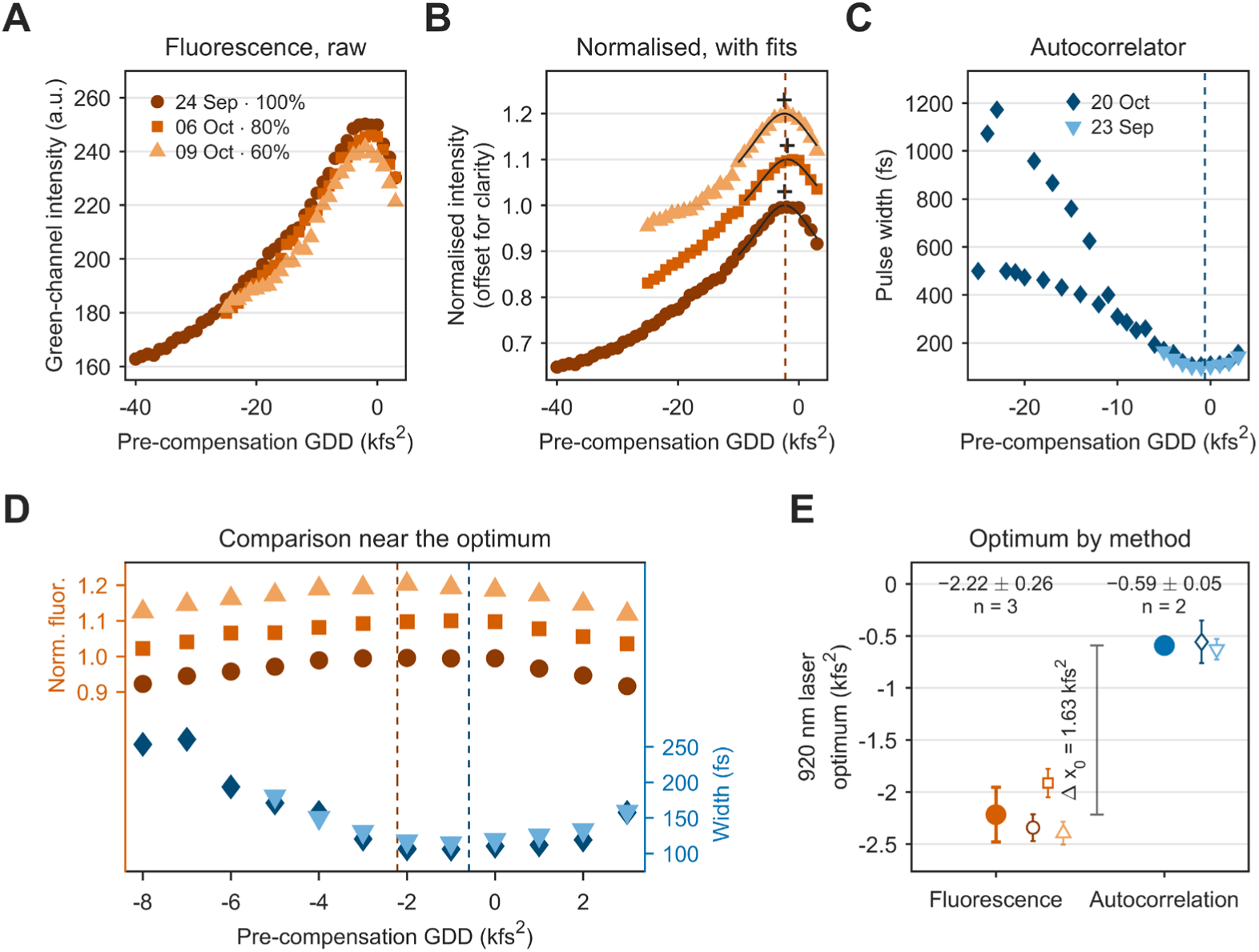
Comparison of fluorescence-based dispersion optimisation with autocorrelation for the G20 nm laser (TOPTICA) with software GDD control. Orange shades, fluorescence; blue shades, autocorrelator; different shades and symbols denote measurement sessions. (A) Raw green-channel fluorescence intensity (a.u.; 8-bit Motion-JPEG video) of 82.8 µM fluorescein in a quartz cuvette versus pre-compensation GDD, for three sweeps recorded on different days and at different laser powers (24 Sep, 100%; 06 Oct, 80%; 09 Oct, 60%). Each point is the maximum pixel value in a 41 × 41 px ROI of the mean of ten frames. This measure differs from the background-corrected ROI mean recommended in the protocol (Fig. 1D). Near the maximum, values approach the 8-bit ceiling of 255; one sweep with detector saturation was excluded. (B) The same sweeps normalised to the maximum of their fit and vertically offset by 0.1 for clarity. Each sweep was fitted (dark lines) with I(x) = B + A/√(1 + ((x − x₀)/w)²), the expected dependence of two-photon fluorescence on GDD for a chirped Gaussian pulse, within ±8 kfs² of the measured maximum. Crosses, drawn slightly above the curves, mark the fitted maxima x₀ (−2.34, −1.91 and −2.40 kfs²); the dashed line marks their mean. Curve fitting was used here to compare the two methods; the protocol itself identifies the optimum as the highest tested mean. (C) Pulse width measured with an autocorrelator in two independent sessions (20 Oct, 23 Sep). Minima were obtained by fitting τ(x) = τ₀√(1 + ((x − x₀)/w)²) within ±6 kfs² of the measured minimum (fits and window boundaries are not shown; all measured points are displayed, but only those within this window contributed to the fit); fitted minima are −0.56 and −0.63 kfs². The dashed line marks their mean. Far from the minimum, where the strongly chirped pulses gave weak second-harmonic signals, the background correction of the autocorrelator software was unstable and the values from 20 October alternated between two levels; these points lie outside the fit window and were not used. (D) Comparison near the optimum: normalised fluorescence (upper part, left axis, offset by 0.1 per series) and pulse width (lower part, right axis, offset by 15 fs per series). Dashed lines mark the mean fitted fluorescence maximum (orange) and the mean fitted pulse-width minimum (blue). (E) Optimal pre-compensation GDD from both methods. Open symbols are individual fits ± SE, with the symbols of (A) and (C). Filled circles are mean ± SD across sessions: −2.22 ± 0.26 kfs² (fluorescence, n = 3) and −0.59 ± 0.05 kfs² (autocorrelation, n = 2; with two sessions, the SD indicates their spread only). The vertical bar marks the difference between the two optima, Δx₀ = 1.63 kfs². Fluorescence was measured at the focus inside the sample and therefore includes the dispersion of every element in the excitation path. For autocorrelation, the beam was re-routed at mirror M1 after the beam expander to the autocorrelator (Fig. 1A), so the elements after M1 (steering mirrors, scanner, scan and tube lenses, objective and the solution at the focus) were not included. The sign of Δx₀ is consistent with the additional positive dispersion of these elements, which was not measured independently. The 1045 nm laser (> 4 W) exceeded the input-power range of the autocorrelator and could not be measured.

The autocorrelation fits, with τ₀ and w fitted independently, gave τ₀ = 98.3 ± 6.1 fs and w = 2.90 ± 0.30 kfs² on 20 October and τ₀ = 97.6 ± 2.5 fs and w = 3.37 ± 0.21 kfs² on 23 September. For a Gaussian pulse these widths correspond to 90 and 97 fs, consistent with the fitted τ₀ within the uncertainties and with the manufacturer’s specification (below 100 fs). The fluorescence sweeps were about twice as broad (w = 6.03 ± 1.23, 6.12 ± 1.53 and 6.48 ± 1.22 kfs²), and only 26–30% of the recorded signal varied with GDD [A/(A + B) of the fit]. Neither the width nor the amplitude A changed systematically between 378, 339 and 228 mW at the sample (A = 71, 64 and 73 counts), whereas a two-photon signal would fall about 2.7-fold over this range if all other conditions were equal. A compressed response of the 8-bit maximum-pixel measure near its ceiling, changed camera settings between sessions or spectral filtering by the steering mirrors (Section 4.3) could contribute; because camera settings were not logged, the present data do not allow the cause to be determined. A fixed, strictly increasing camera response without clipping would preserve the ordering of measured intensities. It does not, however, guarantee an unchanged maximum of a fitted model when the response distorts the curve shape; clipping can also prevent a unique measured maximum from being resolved. The 920 nm sweeps had no reference records, so stability during these sweeps was not assessed; the reproducibility of the optimum rests on the comparison of three sessions.

### 3.4. Heating and fluorescence stability

Under continuous illumination, the samples heated at rates that depended on sample type and power (Fig. 4A,B). On the pollen-grain slide at 480 mW (1045 nm), the temperature jumped at 120 s, coinciding with damage to the sample. On the fluorescent slides, the camera recorded two spots, one from fluorescence and one from reflection at the slide glass, which could not be assigned (Fig. 4C). In the cuvette at 1155 mW, fluorescence at the fixed optimum setting changed over minutes while the temperature rose (Fig. 4E). Slow changes of this kind are what the bracketing references detect without a temperature measurement.

**Fig. 4.**
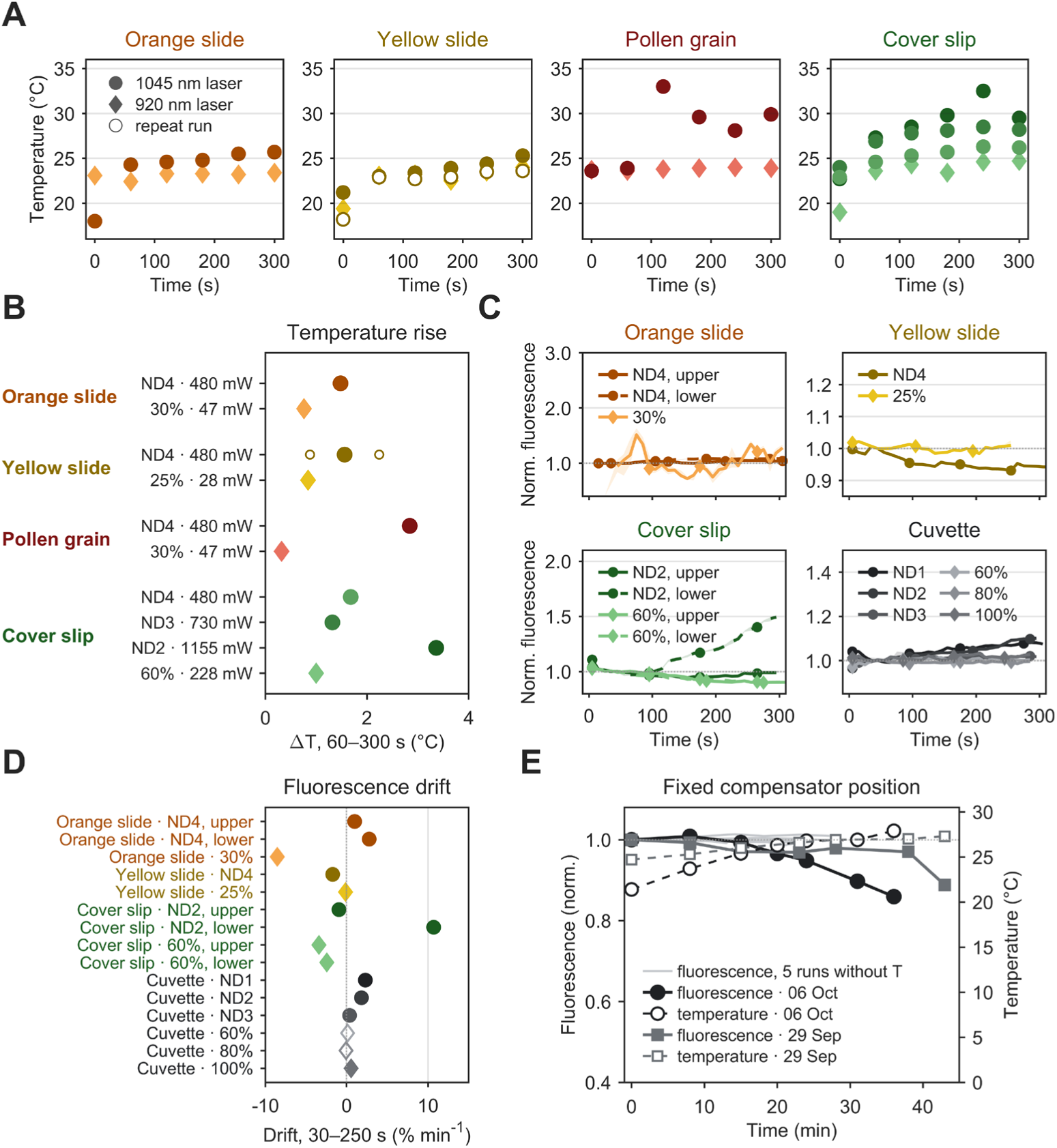
Sample heating and fluorescence stability under continuous illumination. Colour denotes the sample (orange slide, orange; yellow slide, yellow; pollen grain, red; fluorescein under a cover slip, green; fluorescein in a cuvette, graphite). Within each panel, darker shades denote higher power. Circles, 1045 nm laser; diamonds, 920 nm laser. ND1–ND4 denote positions of a neutral-density filter wheel in the 1045 nm beam (ND1, no filter; ND2–ND4, nominal OD 0.5, 1.0 and 2.0). The nominal optical densities apply to visible light; at 1045 nm the attenuation is much weaker, as shown by the measured powers (ND1–ND4: 1440, 1155, 730 and 480 mW); percentages denote the power setting of the 920 nm laser. Powers are average powers at the sample (Thorlabs S425C-L thermal sensor, 10 mm below the objective). **(A)** Temperature of the samples during continuous illumination. Temperature was read every 60 s with a thermal camera and therefore reflects the sample surface rather than the focal volume. The reading at t = 0 s was taken with the laser off, before the sample was brought into contact with the objective. Open symbols, repeat run (yellow slide, 1045 nm laser, ND4; runs on 01 and 02 October). The temperature jump on the pollen-grain slide at 120 s (1045 nm laser, ND4) coincided with damage to the sample (Fig. 5C). Room temperature was approximately 22 °C. **(B)** Temperature rise over 60–300 s (slope of a linear fit × 240 s) for each condition in (A). Yellow slide, ND4: mean of two runs, with small open circles showing the individual runs; all other conditions n = 1. The protocol does not require routine temperature measurement; slow changes during a sweep are detected instead with bracketing reference records. **(C)** Normalised green-channel fluorescence during continuous illumination. Traces show the ROI mean minus the camera offset (8 counts), a 1 s running median and 10 s bins (median, shaded interquartile range), normalised to 30–60 s; symbols mark every eighth bin. Transients exceeding 3 MAD from a 30 s running baseline (±2 s) were masked; they recurred at ∼60 s intervals and coincided with opening the light-shielding curtain for temperature readings, indicating that ambient light reached the camera. On the slides, the camera recorded two light spots, one from sample fluorescence and one from reflection at the slide glass, which could not be assigned; they were analysed separately (upper, solid lines; lower, dashed lines) and are shown to illustrate this difficulty. The protocol excludes reflections from the measurement region. The pollen grain gave no signal above the dark level and is not shown. **(D)** Fluorescence drift over 30–250 s (slope of a linear fit) for each trace in (C). Open symbols, signal < 3 counts above the dark level. **(E)** Fluorescence at a fixed compensator screw position (position 22, the optimum; solid lines, left axis, normalised to t = 0) and sample temperature (dashed lines, open symbols, right axis) for fluorescein in a cuvette with the 1045 nm laser and ND2 (1155 mW), on 06 October and 29 September. Light lines: five runs under the same conditions without temperature recording (24 September, 08 October and 09 October).

### 3.5. Failure modes

Movement of the cuvette during a sweep shifted the apparent maximum from 22 to 14 turns (Fig. 5A, left); with the cuvette fixed, both directions peaked at 22 (Fig. 5A, right). At 730 mW (1045 nm), a pollen-grain slide was damaged within seconds, with violet emission at the focus visible in the selected early-emission frame at 9.2 s and reaching its recorded maximum at 10.9 s (Fig. 5B). At 480 mW, opening the shutter with the beam already focused left damaged regions of 0.28–1.2 mm on pollen-grain, orange and yellow slides; at 730 mW, damage occurred even while the focus was being approached (Fig. 5C).

**Fig. 5.**
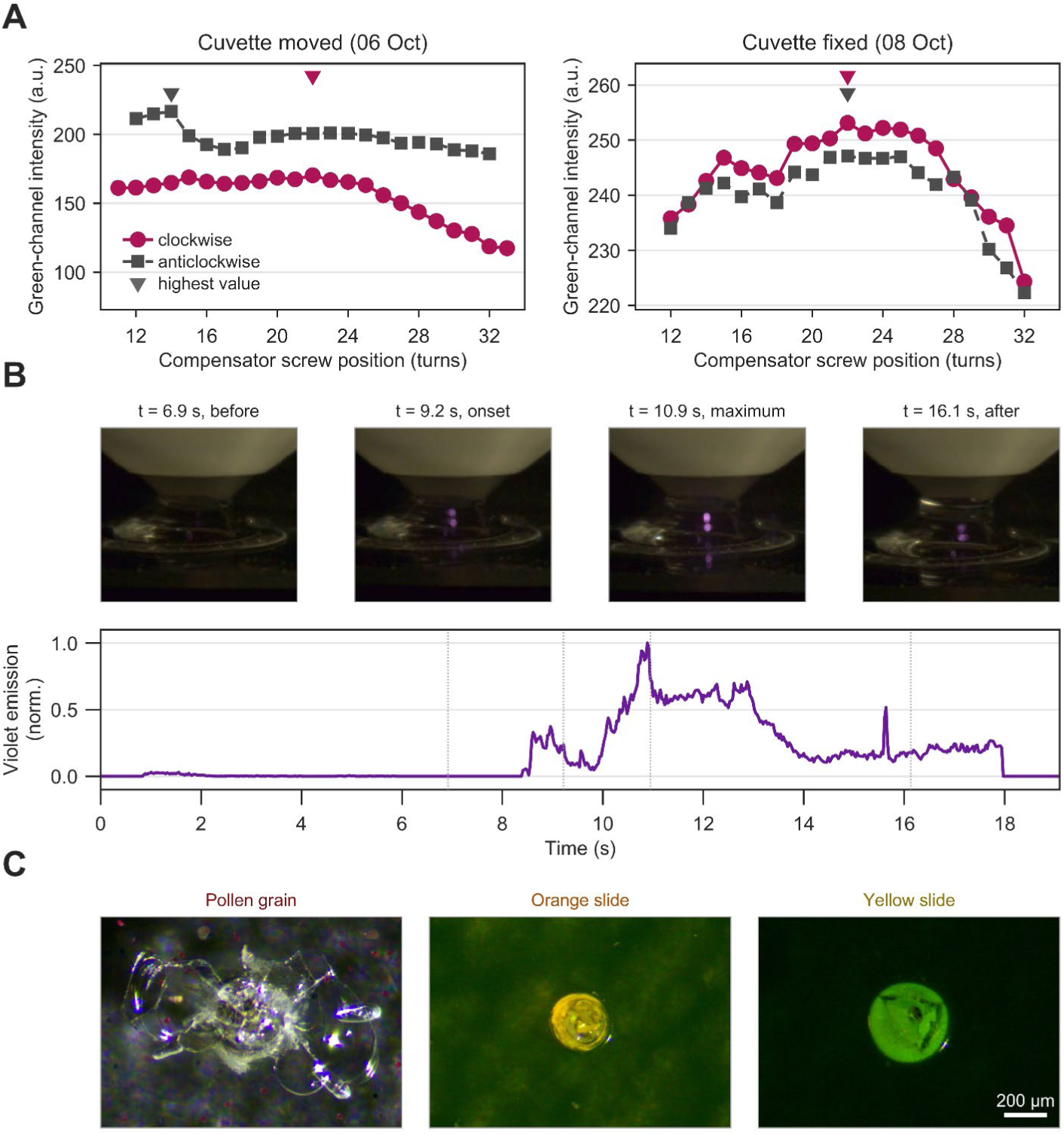
Practical failure modes of fluorescence-based dispersion optimisation. (A) Cuvette displacement during a sweep. Fluorescence versus compensator screw position for clockwise (raspberry, circles, solid lines) and anticlockwise (graphite, squares, dashed lines) sweeps of fluorescein in a cuvette with the 1045 nm laser and ND2. Left: on 06 October, the cuvette moved during the anticlockwise sweep, shifting its highest value from position 22 to 14. Right: on 08 October, the cuvette was fixed and the highest values of both sweeps occurred at position 22 (same data as Fig. 2B,C). Triangles above the data mark the highest value of each sweep. (B) Sample damage at high power. Side-view video frames during illumination of a pollen-grain slide with the 1045 nm laser and ND3 (730 mW), showing localised violet emission accompanying sample damage. Below: violet emission in the region around the focus, computed per frame as the summed (R + B)/2 − G signal above a threshold of 20 and normalised to its maximum. Dotted lines mark the frames shown. The frame labelled “onset” at 9.2 s illustrates early violet emission, rather than a quantitatively determined onset time; the trace begins to rise earlier. Time was derived from the frame rate of the original recording (34.7 frames s⁻¹). (C) Damage left on the pollen-grain, orange and yellow slides after illumination with the 1045 nm laser and ND4 (480 mW). The damage occurred when the shutter was opened with the beam already focused in the sample; with ND3, the samples were damaged even while the focus was being approached. Stereo-microscope images (4× zoom), all cropped to the same field; scale bar, 200 µm. The damaged regions measure approximately 1.2 × 0.7 mm (pollen grain), 0.28 mm (orange slide) and 0.37 mm (yellow slide) across.

## 4. Discussion

### 4.1. What the measurement determines

The procedure finds the compensator setting that maximises the recorded two-photon fluorescence at the sample under the conditions used; it does not measure pulse duration. Like the photomultiplier-based procedure [Lees et al., 2025], it uses fluorescence as a surrogate for peak intensity; with a parked beam and a side-view camera, the microscope’s scanning, collection optics and detector gain play no part. The 1.63 kfs² difference from the autocorrelation optimum shows that the setting found at the sample can differ from one found before the microscope. Its sign agrees with the additional positive dispersion of the optics after M1: the scan and tube lenses alone contain 11.5 mm of N-BK7, about 0.39 kfs² at 920 nm (34.1 fs² mm⁻¹ from the manufacturer’s Sellmeier coefficients), and about 2 mm of water adds 0.03 kfs² [Daimon and Masumura, 2007]. The remainder is consistent with the objective and the steering mirrors but was not measured independently. Within a Gaussian-pulse model with the width from autocorrelation (w ≈ 3.1 kfs²), a difference of 1.63 kfs² would predict a reduction of about 11% relative to the model maximum; this is not a measured signal recovery and assumes that the spectrum at the sample equals that at M1. The present comparison does not independently establish the accuracy of the inferred compensator setting.

### 4.2. Comparison with available approaches

The photomultiplier-based procedure [Lees et al., 2025] uses the microscope’s imaging and detection path. Side-view camera detection provides an alternative, as already demonstrated by Saidi and Shtrahman [Saidi and Shtrahman, 2023], who used a long-working-distance objective and tube lens to image through a custom cuvette window and adjusted GDD using the brightest pixel. Our arrangement uses a compact camera lens and a quartz cuvette with an immersed excitation objective. We develop the sweep workflow around repeated reference records, practical stability warnings and explicit reporting of limitations. The camera also provides a spatial view that helps identify sample displacement (Fig. 5A) and ambiguous light spots (Fig. 4C). Our configuration required hundreds of milliwatts to more than one watt at the sample; higher-NA collection optics, longer exposure or a monochrome camera may reduce the required power, but this was not tested. Users should select the lowest power that provides a usable, unsaturated signal and assess stability relative to the differences between compensator settings. Unlike pulse-characterisation methods [Fittinghoff et al., 1998; Lozovoy et al., 2004; Miranda et al., 2012], the procedure reports a useful compensator setting rather than the temporal pulse profile.

### 4.3. Limitations of the measurement

Interpreting the fluorescence maximum as the shortest pulse requires fixed average power, spectrum and focal geometry as well as excitation far from saturation [Field et al., 2010]; saturation enlarges and flattens the excitation volume [Cianci and Berland, 2004; Nagy et al., 2005]. The same condition underlies the correspondence between the ±5% reference warning level and a 2.5% power change. The quadratic regime can be confirmed with a log–log power series, as done for in vivo retinal imaging, where slopes of 2.05–2.11 were reported [Palczewska et al., 2018].

The recorded sweeps of both lasers were broader than expected from the pulse duration, so the flatness of the maximum describes the recorded signal rather than the underlying fluorescence. For the 1045 nm laser, the maxima spread over 21–24 turns, about 1.7 kfs² assuming a linear compensator (0.58 kfs² per turn); this reflects agreement between sessions with different objectives and sample geometries rather than the precision of the compensator itself. For the 920 nm laser, the fitted maxima varied by 0.26 kfs² (SD); the two measures of spread are not directly comparable, and the fitted width should be reported alongside the optimum.

The UM10-45B steering mirrors were used at 920 nm, outside the manufacturer’s currently specified low-GDD design band of 970–1150 nm for s-polarisation (1020–1080 nm for p-polarisation at 45° incidence, according to the manufacturer’s specification). The spectral response of the installed mirrors at the actual incidence angles and polarisations was not characterised. Spectral filtering and higher-order phase from these coatings are therefore possible contributors to the broader fluorescence sweeps, rather than established explanations. The spectrum after the mirrors was not measured.

During continuous illumination, sample temperature and fluorescence changed together (Fig. 4E), but these data do not isolate heating as the cause of the signal changes; the bracketing references can reveal slow changes, without identifying their cause or excluding a repeatable dependence of power on compensator setting. Pollen-grain and autofluorescent slides, used at low power in the photomultiplier procedure [Lees et al., 2025], were damaged under the conditions used here (Fig. 5B,C); with a camera, a fluorescent solution is preferable, the focus should be approached at low power and the shutter opened only once the focus is set. At 83 µM, fluorescein reabsorbs part of its emission [Sjöback et al., 1995], so the signal depends on the path to the camera-facing wall, and cuvette movement shifted the apparent maximum (Fig. 5A). A water-only control is recommended to exclude laser leakage and hyper-Rayleigh scattering [Xu et al., 1997].

The measurements reported here were recorded as 8-bit Motion-JPEG video and served to derive the quality checks; the protocol builds on them by prescribing lossless recording, logged camera settings, a log–log power series to confirm the quadratic regime and, at 1045 nm, a water-only control. Only GDD is optimised: for pulses longer than about 50 fs, GDD dominates pulse broadening in the microscope [Field et al., 2010], but higher-order phase [Xi et al., 2009] and energy outside the main peak of the pulse [Chong and Török, 2024] cannot be corrected by adjusting GDD. The autocorrelation was measured before the scan optics, and the measured pulse duration (98 fs) lies near the lower limit of the autocorrelator’s range (70 fs). We tested two lasers, one camera and one principal cuvette configuration, with small numbers of sessions.

### 4.4. Possible applications

The approach may be useful in other systems with adjustable compensation, side-view access to the focus and a sufficiently strong, unsaturated signal from a suitable fluorescent reference. In neuroscience, two-photon microscopes are often custom-built or modified [Janiak et al., 2022] and reconfigured between experiments, and heating limits the usable power in tissue [Podgorski and Ranganathan, 2016; Schmidt and Oheim, 2020]; checking the setting after each change of objective, wavelength or beam path is a natural use. Water adds 12.6 fs² mm⁻¹ of dispersion at 920 nm and close to zero at 1045 nm [Daimon and Masumura, 2007; Coello et al., 2007], which suggests a small contribution from water over a few millimetres compared with the spread observed here. This does not establish transfer of the cuvette setting to tissue: differences in coverslips, immersion, objective correction and sample optics may also matter, and transfer was not tested. Other candidate uses are two-photon photostimulation paths [Lees et al., 2024] and two-photon polymerisation, whose initiation also depends on two-photon absorption [Samsonas et al., 2023]; there, a cover-slip-bottomed cell and a dye without one-photon absorption at the excitation wavelength would be needed. For depth-representative measurements, a fluorophore embedded in a tissue-mimicking or nanodiamond phantom [Janiak et al., 2026; Wróbel et al., 2016; Listewnik et al., 2021] could replace the cuvette.

## 5. Conclusion

Side-view imaging of two-photon fluorescence provides a practical way to adjust dispersion compensation at the sample independently of microscope scanning and detection. Building on established side-view compensation methods, we describe a sweep workflow with reference records and practical stability warnings informed by observed failure modes. The recorded maxima show agreement across sessions, with precision limited by instability, camera response and proximity to clipping. For a 920 nm fibre laser, the mean fitted fluorescence maximum differed by 1.63 kfs² from the mean autocorrelation minimum measured before the scan optics, illustrating that the selected setting depends on the optical path included in the measurement. Open software and a verification dataset support reproducible analysis. Photostimulation and polymerisation paths are potential applications where side-view access and a suitable fluorescent reference are available; these extensions were not tested here.

## 2. Methods

### 2.1. Lasers and excitation path

Two femtosecond lasers were used (Fig. 1A). A Menlo Systems YLMO (1045 nm, 100 MHz; manufacturer’s test report: 75 fs assuming a sech² pulse shape, 4.1 W, p-polarised) has a user-settable pre-chirp compressor covering 0 to −30 000 fs² over 52 turns of an adjustment screw, about 0.58 kfs² per turn assuming a linear dependence. A TOPTICA FemtoFiber ultra 920 (920 nm, 80 MHz; manufacturer’s specification: pulse duration below 100 fs, average power above 1.5 W) has motorised GDD pre-compensation adjustable in the laser software from −40 000 to +3000 fs²; negative settings add negative GDD that pre-compensates the positive dispersion of the optics downstream. Its power setting was 60–100% of maximum.

Each laser had its own 2× beam expander of two N-BK7 plano-convex lenses (Thorlabs LA1131-B, f = 50 mm, and LA1509-B, f = 100 mm; AR 650–1050 nm) and a shutter (Thorlabs SH05R/M with SC10 controller). A manually translated mirror selected the laser, transmitting the 1045 nm beam and reflecting the 920 nm beam, and three low-GDD mirrors (Thorlabs UM10-45B) steered it to an X–Y galvanometric scan module (Sutter SA-GEN-RLYD-GG-SCNBX), which was parked during the measurements. A scan lens (Thorlabs LA1131-B, f = 50 mm, Ø1″) and a tube lens (LA1979-B, f = 200 mm, Ø2″) relayed the beam to the objective, and a mirror (Thorlabs UME1-AG) directed it downwards to an Olympus XLUMPlanFL 20×/1.00 W water-dipping objective. One 1045 nm sweep (14 May) used a Zeiss W Plan-Apochromat 20×/1.0 DIC VIS-IR water-dipping objective. In the 1045 nm beam, a filter wheel with absorptive neutral-density filters (Thorlabs FW212CNEB) provided four positions (ND1, no filter; ND2–ND4, nominal OD 0.5, 1.0 and 2.0). Because these optical densities are specified for visible light, the attenuation at 1045 nm was much weaker, and the measured average powers at the sample were 1440, 1155, 730 and 480 mW. For the 920 nm laser, power settings of 25, 30, 60, 80 and 100% gave 28, 47, 228, 339 and 378 mW at the sample. Power was measured with a thermal sensor (Thorlabs S425C-L, read out with a PM100D2 console) placed 10 mm below the objective and cross-checked with a photodiode sensor (Thorlabs S121C), which read 9–22% higher (median 13%) where both were used.

### 2.2. Reference samples

Fluorescein free acid (Sigma-Aldrich 46955; 332.31 g/mol) was dissolved with triethylamine in deionised water to a 1.655 mM stock and diluted to 82.8 µM (pH 9.0); the recipe is in Supplement 1. The solution filled a 10 mm quartz cuvette (Hellma QS) to the rim, giving a flat liquid surface, and the objective was immersed without touching the walls of the cuvette. Heating and stability measurements also used orange and yellow autofluorescent plastic slides (Chroma 92001) and a pollen-grain slide (Carolina 304264), the reference samples recommended in [Lees et al., 2025], and a drop of fluorescein (50–100 µL) between a microscope slide (1.0–1.15 mm thick) and a cover slip (0.13–0.16 mm).

### 2.3. Fluorescence detection

A colour CMOS camera (Thorlabs CS165CU, controlled with ThorCam) with a 3.5 mm f/1.4 lens (Thorlabs MVL4WA, aperture fully open) viewed the focus from about 5 cm at 90° to the excitation axis through a flat cuvette wall. No emission filters were used; the camera has a built-in, removable IR-blocking filter, which was kept in place. All recordings except the sample-damage sequence in Fig. 5B were 8-bit Motion-JPEG video. Camera exposure, gain and gamma were not logged for these recordings; amplitudes are therefore compared only within a sweep. Motion-JPEG stores colour as luminance with sub-sampled chrominance, so its green channel is reconstructed rather than read directly from green sensor sites, and its 8-bit values need not be linear in intensity. The protocol therefore prescribes lossless TIFF stacks (Supplement 1).

### 2.4. Acquisition and quality checks

In this study, the compensator was moved to a preload position before each record and approached the target setting from a fixed side to remove backlash; after at least 5 s of settling with the shutter closed, the shutter was opened and, after a 1 s pre-exposure, a short video was recorded. Each record consisted of ten consecutive video frames (about 0.29 s at 34.7 frames s⁻¹). Test settings changed monotonically, and clockwise and anticlockwise sweeps were analysed separately. Reference records were acquired in the 1045 nm sweeps; the 920 nm sweeps were single-direction sweeps without reference records, one per session. For the 1045 nm laser, 13 sweeps were acquired over nine sessions; for the 920 nm laser, four sweeps were acquired over four sessions, one of which reached the 8-bit ceiling and was excluded.

The recommended protocol (Fig. 1C; Supplement 1) uses the same preload, settling and pre-exposure, but acquires ten frames at a 1.0 s frame-start interval in a lossless format with logged camera settings, repeats a reference record after at most five test records, and follows a coarse sweep with at least three fine sweeps per direction.

Table 1 summarises practical quality checks informed by the failure modes observed during method development. The 5% thresholds for temporal variability, within-record drift and reference stability were selected by the authors as operational warning levels; they were not independently validated as limits on the accuracy of the compensator setting. Exceeding these thresholds flags a sweep for closer inspection rather than automatically invalidating its observed maximum. Interpretation should consider the magnitude and time course of the instability relative to the signal differences between settings near the maximum, together with agreement between repeated sweeps. Clipping, sample displacement or a maximum at the boundary of the tested range can prevent an optimum from being resolved and must be reported explicitly.

### 2.5. Image analysis and software

Figures 2A,B,D and 3 were analysed using the maximum green-channel pixel value of the ten-frame mean image (Fig. 2) or within a 41 × 41 px ROI of that mean image (Fig. 3), measured in Fiji (ImageJ) [Schindelin et al., 2012]. Figure 2C additionally evaluates a fixed signal ROI mean minus a fixed background ROI mean for each frame, summarised as mean ± SD over ten frames. These images were exported from Motion-JPEG video to TIFF; this conversion does not restore sensor linearity or information lost during compression. For prospective acquisition, the protocol recommends the background-corrected ROI mean using green Bayer sites from native colour-sensor data, the green channel from verified linear RGB data, or all pixels from monochrome data (Fig. 1D).

The positions in Fig. 2A are measured maxima, without fitting. Maximum-pixel sweeps (Figs 2B and 5A and the thin lines in Fig. 2C) are shown as recorded or normalised to their own maximum, without baseline correction or smoothing. For the comparison in Fig. 2C, the recordings of 8 October were exported as ten-frame TIFF stacks and analysed with the protocol software. All figures were produced with MATLAB R2020a.

The analysis software (analyse_dispersion_v10; MATLAB R2020b or later with Image Processing Toolbox, and Python 3.8 or later with NumPy, tifffile and Matplotlib) reads the TIFF stacks and a five-column metadata file, applies the quality checks in Table 1 and reports a candidate optimum. Both implementations were tested against a synthetic verification dataset of ten cases with expected values derived from the construction of the data, and both reproduced all expected records, frames and warning flags. The analysis code, the verification dataset and the source data of the figures are available in the MOST Wiedzy Open Research Data catalogue of Gdańsk University of Technology [Czubek et al., 2026].

### 2.6. Curve fitting

To compare fluorescence with autocorrelation (Fig. 3), fluorescence sweeps were fitted within ±8 kfs² of the measured maximum with I(x) = B + A/√(1 + ((x − x₀)/w)²), the dependence expected for a chirped Gaussian pulse, and pulse-width sweeps within ±6 kfs² of the minimum with τ(x) = τ₀√(1 + ((x − x₀)/w)²). Both were unweighted least-squares fits (MATLAB `fminsearch`, residuals scaled by the largest measured value, refined by a second run from the first solution). Repeated measurements at the same setting were averaged before fitting, and sweeps reaching the 8-bit ceiling (any value ≥ 254.5) were excluded. Uncertainties of fitted parameters are standard errors from the numerical Jacobian of the fit, and between-session values are given as mean ± SD. For a Gaussian pulse, w = τ₀²/(4 ln 2), so the width of a fluorescence sweep can be compared with the pulse duration from autocorrelation, and the signal lost at a setting Δx from the optimum is 1 − 1/√(1 + (Δx/w)²). The protocol itself does not fit; it reports the highest tested mean of each sweep together with its quality flags.

### 2.7. Autocorrelation

For the 920 nm laser, the beam was diverted at mirror M1 after the beam expander to a single-shot second-harmonic intensity autocorrelator (Femto Easy ROC-PS3; pulse-duration range 70 fs–3 ps; phase-matching option for 920 and 1045 nm; integrated afocal beam reducer; 12-bit CMOS detection), in which two copies of the beam cross in a nonlinear crystal and the second-harmonic trace is imaged onto the camera. Pulse durations were obtained by the manufacturer’s software assuming a Gaussian pulse shape (deconvolution factor 1.414). Measurements were made in two sessions (23 September and 20 October 2026). The 1045 nm laser exceeded the input-power range of the autocorrelator (3.5 W average power) and was not measured.

### 2.8. Temperature and fluorescence stability

Sample temperature was read every 60 s with a thermal camera (FLIR i7; emissivity set to 0.95, factory calibration), so it reflects the sample surface rather than the focal volume. Because one emissivity value was used for all samples, absolute temperatures are approximate; the analysis relies on temperature changes over time. The temperature rise (Fig. 4B) is the slope of a linear fit over 60–300 s multiplied by 240 s; repeated runs under the same conditions were averaged. A temperature jump on the pollen-grain slide at 120 s was checked and kept as a valid reading.

For the stability traces (Fig. 4C), the ROI mean of each video frame was corrected for the camera offset (8 counts) and smoothed with a 1 s running median. Transients were detected as deviations of more than 3 scaled median absolute deviations from a 30 s running median and masked for ±2 s; they coincided with opening the light-shielding curtain for temperature readings. The masked trace was summarised in 10 s bins (median and interquartile range) and normalised to the median of the bins at 30–60 s. Drift (Fig. 4D) is the slope of a linear fit over 30–250 s in % per minute; traces whose median signal over 30–250 s was less than 3 counts above the dark level are marked separately.

### 2.9. Sample damage

Sample damage was recorded from the side in an uncompressed video (34.7 frames s⁻¹). Violet emission was quantified in a fixed region around the focus as the per-frame sum of (R + B)/2 − G above a threshold of 20 (8-bit values) and normalised to its maximum. Damaged samples were imaged afterwards with a trinocular stereo microscope (AmScope SM-1TS-L6W-M) equipped with a 0.37× camera adapter (AmScope RU037) at 4× zoom setting, with the scale calibrated for each zoom setting with a stage micrometer standard (Bresser 12897); the images were cropped to the same 720 × 504 px field (1.96 µm per pixel), and the size of the damaged regions was measured on the calibrated images.

### 2.10. Statistics

Values are given as mean ± SD. For a single record, the SD is taken over its ten frames (Fig. 2C); for the 920 nm laser, over sessions, with each fluorescence sweep and each autocorrelation measurement acquired in a separate session (n = 3 and n = 2; Fig. 3E). With n = 2, the SD only describes the spread of the two values. Uncertainties of fitted parameters are standard errors from the fit (Section 2.6). The agreement between the protocol metric and the maximum-pixel measure (Fig. 2C) is given as the Pearson correlation coefficient.

## Supporting information

Supplement1_Protocol

## Data availability

Data underlying the results presented in this paper (the source data of all figures), the analysis code (MATLAB and Python, version 10), the synthetic verification dataset and the laboratory protocol are available in the MOST Wiedzy Open Research Data catalogue of Gdańsk University of Technology [Czubek et al., 2026]. The original video recordings are available from the authors upon reasonable request.

## Supplemental document

See Supplement 1 for supporting content.

