## Supplement1_Protocol for "Optimisation of femtosecond dispersion compensation at the sample using side-view two-photon fluorescence imaging"

**Supplement 1**

*Optimisation of femtosecond dispersion compensation at the sample using side-view two-photon fluorescence imaging*
Jakub Czubek, Grzegorz Orlicki, Jacek Łubiński, Małgorzata Szczerska and Filip Janiak
Data, analysis software and verification dataset: https://doi.org/10.34808/em3y-6b44

*Laboratory protocol, version 14, 23 November 2026*

### Optimising femtosecond laser dispersion compensation using fluorescence

#### 1 Purpose

Use two-photon fluorescence to find the optimum dispersion compensation at the intended focal plane, with the microscope excitation optics and objective in place. The procedure applies to any femtosecond laser system with adjustable internal or external dispersion compensation. An autocorrelator is not required.

The result is the compensator setting that maximises two-photon fluorescence under the recorded conditions. It does not measure pulse duration, demonstrate zero residual dispersion or determine the sign or magnitude of chirp. A pulse-duration claim requires an independent measurement, such as autocorrelation at the sample plane.

Keep the sample, collection geometry, source output, attenuation and camera settings fixed while changing the compensator setting. Measure fluorescence, identify the maximum and refine the setting with smaller steps.

#### 2 Equipment and software

| Item | Requirement |
|---|---|
| Excitation system | Femtosecond laser, adjustable dispersion compensation, shutter and the objective and optics used for the intended experiment. |
| Camera | Any monochrome or colour camera suitable for recording the fluorescence, with manual acquisition settings, suitable collection optics and a stable mount. |
| Reference sample | A stable fluorescent preparation suited to the wavelength and mounting arrangement. Section 3 gives a fluorescein preparation used in our experiments. |
| Sample support | A slide, coverslip, cuvette, chamber or agarose preparation, selected for the experiment and objective (Section 4). |
| Solution preparation | If preparing the stock in Section 3: fluorescein free acid, deionised water, triethylamine, a laboratory balance, pipettes and a 25 mL volumetric flask. |
| Software | Acquisition software compatible with the camera. Fiji is free and open source (https://fiji.sc/). Optional MATLAB and Python analysis files are described in Appendices A and B. |

Tested example equipment: Menlo Systems YLMO and TOPTICA FemtoFiber ultra 920 lasers; Thorlabs CS165CU colour camera controlled with ThorCam. These identify the equipment used for the experiments; other compatible lasers and cameras may be used.

Follow the local laser operating procedure. Handle triethylamine in a fume cupboard under the applicable chemical assessment.

### 3 Prepare the reference sample

**The method does not require the exact fluorescein concentration specified below.**
The recipe describes the solution used in our experiments, approximately 83 µM, and provides a reproducible starting point. Adjust concentration to obtain clear, stable fluorescence within the camera's linear range. Select the solution composition and acquisition settings before each sweep and keep them unchanged throughout acquisition.

#### 3.1 Prepare the stock solution

**1.** Use fluorescein free acid, CAS 2321-07-5, molecular mass 332.31 g/mol [1]. If using a different chemical form, recalculate the mass for the intended molar concentration.

**2.** Weigh 13.75 mg of fluorescein free acid into a suitable vessel. Add approximately 20 mL of deionised water and 62.5 µL of triethylamine. Stir for 15 min and confirm complete dissolution.

**3.** Transfer the solution into a 25.00 mL volumetric flask. Rinse the original vessel with small portions of deionised water and add all rinses to the flask, keeping the liquid below the calibration mark. Allow the solution to reach room temperature, then add water to the mark and mix. The final volume, including all rinses, is 25.00 mL. This gives a nominal stock concentration of 1.655 mM.

#### 3.2 Prepare the working solution

**4.** Mix 125 µL of stock with 2.375 mL of deionised water to give 2.500 mL at approximately 83 µM. Alternatively, mix 250 µL of stock with 4.750 mL of water to give 5.000 mL at the same concentration. Mix thoroughly.

**5.** Prepare enough solution for the chosen holder and immersion arrangement. Use one batch throughout a sweep. Record the recipe or stock dilution and, when measured, pH. Protect the solution from light between measurements.

### 4 Arrange the sample and camera

**Choose a sample configuration representative of the intended experiment.**
Options include a fluorescent reference with a water-immersion meniscus, a microscope slide or coverslip preparation, a cuvette containing fluorescent solution, or a stable fluorescent reference in an agarose preparation representative of zebrafish mounting. Use the intended objective and immersion medium, including the relevant glass, water and agarose layers in the excitation path.

**1.** For a slide, coverslip or agarose preparation, place a stable fluorescent reference at the intended imaging depth. Select and secure the preparation before setting the camera.

**2.** Secure the preparation. Set the reference at a representative imaging depth and establish the required immersion contact. Keep the objective clear of the holder and glass.

**3.** Park the excitation focus at a fixed position. Position and focus the camera so that the fluorescent reference is visible. Exclude bubbles, meniscus reflections and bright holder surfaces from the measurement region.

**4.** Reduce ambient light and use suitable emission and excitation-blocking filters. Fix the camera, objective, sample and collection optics for the complete sweep.

##### 4.1 Example cuvette arrangement

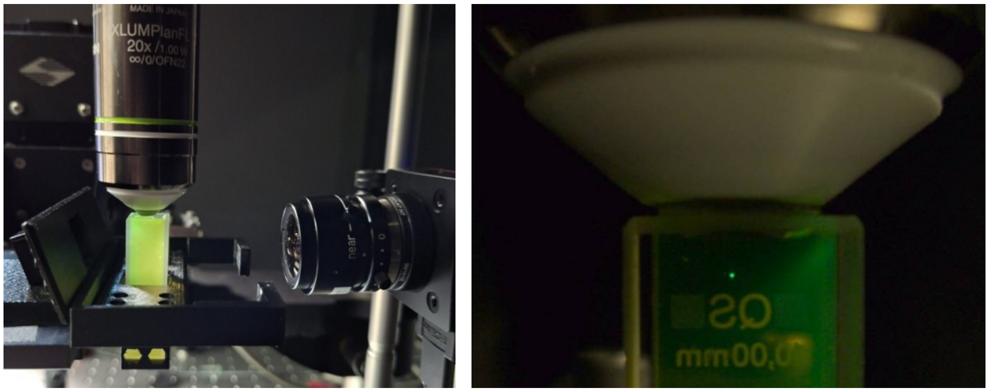


*Figure S1. Example cuvette configuration. Left: objective, open cuvette and side-view camera under room illumination. Right: localised fluorescence inside the cuvette. Photographed objective: Olympus XLUMPlanFL 20×/1.00 W, water-dipping.*

For this arrangement, fill the cuvette to the rim so that the liquid surface is flat, and immerse the dipping objective in the solution without touching the walls of the cuvette. Position the focus away from walls and the surface. View through a flat side wall, approximately perpendicular to the excitation axis. Note the focal depth and camera-facing wall clearance; keep both fixed during the sweep. At approximately 83 µM the solution absorbs part of the green emission, so the detected signal depends on the path from the focus to the camera-facing wall. Keep this distance fixed within and between compared sweeps.

### 5 Configure the camera and run a pilot

#### 5.1 Select the image representation

Use monochrome intensity images or a suitable channel from linear RGB images; for fluorescein, use the green channel. Native Bayer data are an additional option for colour cameras. Keep the chosen representation unchanged throughout each sweep.

**What is a Bayer TIFF?**
A Bayer TIFF stores raw colour-camera data before reconstruction into RGB. Each pixel records one intensity through a red, green or blue filter; the usual repeating pattern contains two green pixels, one red and one blue. Reconstruction into RGB is called demosaicing. Analyse native Bayer data using the correct filter pattern, such as RGGB or BGGR. A single-channel TIFF may instead be monochrome. Bayer acquisition is optional: monochrome and linear RGB images can also be used.

#### 5.2 Fix acquisition settings

**1.** Disable automatic exposure, gain and white balance where applicable. Use a linear intensity output; disable gamma adjustment, adaptive enhancement and frame averaging. Record exposure, gain, gamma and image mode, or save the camera profile; without these records, signal amplitudes cannot be compared between sessions.

**2.** Use lossless images. For the supplied scripts, save each measurement as a TIFF stack containing exactly ten separate frames. Preserve native pixel values when exporting or converting files in Fiji [2].

**3.** Set a nominal frame-start interval of 1.0 s, with exposure and readout fitting inside that interval. Check the actual timing in the acquisition log. Record the interval used.

**4.** Determine the clipping code of the stored data from the acquisition settings or camera documentation. A 16-bit container alone does not establish the effective camera ceiling. Keep at least 5% headroom at the brightest setting.

#### 5.3 Run a pilot

**5.** Warm up the laser according to its operating instructions. Choose source output, attenuation and exposure that give a clear signal across the sweep without saturation. Keep these settings fixed.

**6.** At low illumination, check that background-corrected signal increases proportionally with exposure before approaching saturation. Then acquire three ten-frame records at one reference setting using the illumination schedule in Section 6. Check signal stability using Section 8.

### 6 Acquire the compensator sweep

#### 6.1 Define the sweep

Choose a safe setting range, coarse step size, adjustment direction and reference setting. Start with the reference, repeat it after no more than five test points and finish with the reference. Record the actual control units. Keep each directional sweep and its repeats in separate folders.

For a mechanical adjuster, choose a safe preload greater than the backlash. Reach every test and reference setting from the same final direction, including the first test after returning to a reference. Use target minus preload for increasing readout, or target plus preload for decreasing readout. Apply this only where the control requires backlash compensation; remain within safe travel.

For open liquid preparations, define an acceptable liquid-level change in millimetres before starting. Check the level and immersion bridge at each reference and at the end. Restore the preparation and restart if the tolerance is exceeded; avoid topping up during a sweep.

#### 6.2 Acquire each record

**1.** Close the shutter and set the compensator. Apply the selected final approach direction where relevant.

**2.** Allow at least 5 s for settling with the shutter closed. Use the same settling and illumination schedule for every record.

**3.** Open the shutter. After a fixed 1 s pre-exposure, acquire ten individual frames at the selected interval. Close the shutter after the final exposure.

**4.** Record the filename, compensator setting and first-frame time. Check the stack for saturation, movement, reflections and unchanged settings.

If available, observe the software laser-status or power indication for unexpected changes. An existing beamsplitter and photodiode monitor may be used. Separate power-meter readings at every point and routine temperature measurements are not required. Bracketing references provide an indirect check of power stability. With other conditions unchanged, two-photon fluorescence scales with the square of power, so the ±5% reference warning level corresponds to a power change of about 2.5%.

#### 6.3 Refine and repeat

**5.** Complete the coarse sweep, identify the brightest region and repeat with smaller steps on both sides of the maximum. If the highest point is at a range boundary, extend the safe range. Use the same reference setting and acquisition schedule for the fine sweeps.

**6.** Repeat the fine sweep to check reproducibility. For numerical reporting of the optimum, at least three independent fine sweeps per reported direction are recommended. Keep clockwise and anticlockwise results separate where relevant.

### 7 Measure fluorescence and select the optimum

#### 7.1 Select fixed regions of interest

**1.** Inspect the endpoints and brightest unsaturated images. Choose a signal ROI containing the fluorescence throughout the sweep, with a margin for small movement. Exclude reflections, walls and the meniscus.

**2.** Choose a separate nearby background ROI outside the fluorescence. Use an uncontaminated region at least as large as the signal ROI where possible. Keep both ROIs fixed for the complete sweep.

#### 7.2 Measure in Fiji or use the supplied scripts

**3.** For monochrome TIFF stacks, measure the recorded intensity. For RGB stacks, use Image > Color > Split Channels and measure the green stack; measuring the RGB stack directly averages all three channels, with weighting depending on the conversion settings. For native Bayer TIFFs, use the Bayer mode of the supplied scripts to select green sensor sites.

**4.** In Fiji, open Analyze > Set Measurements and select Mean gray value. Add the signal and background rectangles to the ROI Manager, select both, and use More > Multi Measure. Measure all slices with one row per slice [3]. Save the ROI set and results; use the same ROIs for every stack.

**5.** For each frame, subtract the background ROI mean from the signal ROI mean. For each compensator setting, calculate the mean and sample standard deviation of the ten corrected frame values. Plot the corrected mean against the actual setting, with temporal standard deviation as error bars.

The optional MATLAB and Python files perform these measurements and report the warnings in Section 8. They support monochrome, linear RGB and native Bayer TIFF stacks. Appendices A and B give the five-column metadata format and commands.

#### 7.3 Select and confirm the setting

**6.** Review Section 8 and identify the highest corrected mean. If the highest mean lies at a range boundary, or if the clipping code is reached at or next to it, the optimum is not resolved; report this explicitly. Refine around that region until the setting is resolved at the required step size. If neighbouring settings remain indistinguishable, retain the corresponding interval.

**7.** Confirm the setting in independent fine sweeps. Record each optimum, the step size and adjustment direction. For numerical reporting, calculate the mean and standard deviation of the independent optimum settings.

**8.** Save the original images, settings, ROI definitions, point log or CSV, analysis results and code version. Keep excluded data with a short reason for exclusion.

#### 8 Check the measurement

Review these checks for every sweep. The 5% values for frame variability, within-record drift and reference stability are author-selected warning levels, also used by the supplied scripts; they were not independently validated as limits on the accuracy of the compensator setting. Exceeding them flags a sweep for closer inspection rather than automatically invalidating its maximum. The 95% headroom is a separate recommendation that protects against clipping.

| **Check** | **Recommended condition or warning level** |
|---|---|
| Acquisition | Ten individual frames per record at a 1.0 s frame-start interval; unchanged camera settings, image mode and scaling; exposure, gain and gamma recorded, automatic settings off. |
| Digital range | No clipped fluorescence; maximum in either ROI no greater than 95% of the verified clipping code (recommended headroom). Clipping at or next to the highest mean leaves the optimum unresolved. Check original data for clipping hidden by processing. |
| Frame variability | Positive corrected mean; temporal coefficient of variation no greater than 5% for each record (author-selected warning level). |
| Within-record drift | Difference between the final three-frame mean and first three-frame mean, divided by the ten-frame mean: absolute value no greater than 5% (author-selected warning level). |
| Reference stability | Every reference within ±5% of the first reference (author-selected warning level). References bracket the sweep; no more than five test points between references are recommended. |
| Preparation | Fixed valid ROIs and geometry; stable immersion contact; no bubbles, moving reflections or excessive liquid-level change. |

If a warning occurs, inspect the individual frames and the reference trace. Judge the magnitude and time course of the instability against the signal differences between settings near the maximum, and compare repeated sweeps. Where the cause can be corrected, correct it and repeat the complete sweep. A rising reference signal suggests heating or source drift; a falling signal suggests bleaching, evaporation, power loss or movement. Allow the system to stabilise, replace the solution if needed, and do not interpret a trend that follows acquisition order as a compensation effect. Keep the ROIs fixed: if the fluorescence leaves them, restore the geometry and repeat rather than moving the ROIs between records. Do not combine favourable records from different sweeps. If the maximum is at an endpoint, extend the safe range; if the signal is insufficient, improve collection or exposure during a new pilot. If the maximum shifts with adjustment direction, check backlash and report each direction separately.

#### 9 Short measurement record

Complete once per configuration. Attach the camera profile, ROI set and point log or metadata CSV.

| Field | Entry |
|---|---|
| Run | Date / operator / folder: ____________________________________ |
| Laser and optics | Laser / wavelength: ___________________ Objective: ___________________ |
| Sample and geometry | Preparation / batch / mount: ____________________________________<br>Immersion / focal depth / relevant wall clearance: ___________________ |
| Camera | Model / image mode: _______________ Exposure / gain / gamma: ___________<br>Clipping code: _________ Bayer pattern (if applicable): ___________<br>Emission / excitation-blocking filters: _________________________ |
| Sweep | Range / step / unit: ___________________ Direction / reference: ___________ |
| Timing | Frame interval: ____ s Settling: ____ s Pre-exposure: ____ s |
| Result and notes | Best setting / repeats: ___________________<br>Files, checks or deviations: ____________________________________ |

### 10 Software and reagent links

[1] Fluorescein free acid: Sigma-Aldrich 46955

[2] Fiji, free and open-source image analysis: https://fiji.sc/

[3] ImageJ ROI Manager and Multi Measure: ImageJ User Guide

[4] MATLAB ROI selection: drawrectangle documentation

[5] Python packages: NumPy; tifffile; Matplotlib

### Appendix A MATLAB analysis file

Use the separate file analyse_dispersion_v10.m with MATLAB R2020b or later and Image Processing Toolbox [4]. Both supplied scripts accept 8-bit or 16-bit integer TIFF stacks. The source code is supplied as a separate file.

#### A.1 Prepare the script metadata

For either script, place the following CSV beside the TIFF stacks. Use one row per ten-frame record, in acquisition order. Only these five fields are required. Additional columns, such as power or temperature, are kept in the output but not checked.

| Column | Content |
|---|---|
| File | TIFF filename or relative path, including extension; unique for each record. |
| Kind | test or reference. |
| Setting | Actual numeric compensator setting. |
| Unit | A consistent unit label, such as turn, fs2 or mm. |
| Time_s | Elapsed time of the first frame, in seconds. |

File,Kind,Setting,Unit,Time_s

#### A.2 Run MATLAB

**1.** Save the file as analyse_dispersion_v10.m, add its folder to the MATLAB path and run analyse_dispersion_v10. Select the CSV, choose the verified clipping code from the list (255, 1023, 4095, 16383 or 65535; choose Other for another verified value) and choose Monochrome, Linear RGB or Native Bayer. Enter the Bayer pattern only for Bayer input.

**2.** Select a bright unsaturated preview stack. Zoom as needed, leave zoom or pan mode, then draw the red Signal ROI and double-click inside it to confirm. Repeat for the blue Background ROI. Instructions for each step appear in the figure title and the Command Window. Confirmed ROIs are locked.

**3.** Review records_v10.csv, frames_v10.csv, roi_v10.png, sweep_v10.png and status_v10.txt in the timestamped analysis_v10 folder. Check the listed warnings, the reference trace and whether the optimum is resolved; confirm the candidate with a finer sweep and repeats.

SHA-256: 8c23b64654a60fedd5e3d4479542506b2ca5fa110fe9e937382cbc02ef7c2196

### Appendix B Python analysis file

Use the separate file analyse_dispersion_v10.py with Python 3.8 or later (tested with Python 3.12) and NumPy, tifffile and Matplotlib [5]. Use the same TIFF stacks and five-column CSV described in Appendix A.1.

#### B.1 Install and run

**1.** Install the packages in the Python environment used for analysis:

```
python -m pip install numpy tifffile matplotlib
```

If the TIFF compression requires an external decoder, such as LZW, also install imagecodecs. Interactive ROI selection requires a graphical Matplotlib backend.

**2.** Run the following command for monochrome TIFFs. Replace CEILING and the filenames with the values used for the dataset:

```
python analyse_dispersion_v10.py sweep.csv --ceiling CEILING --mode mono --preview ref.tif
```

**3.** For linear RGB, use --mode rgb. For native Bayer, use --mode bayer and add --pattern RGGB, BGGR, GRBG or GBRG to match the stored layout. A clipping code other than 255, 1023, 4095, 16383 or 65535 triggers a reminder to verify it on a saturated test image.

**4.** Click two opposite corners of the signal ROI, then two corners of the background ROI. To enter fixed ROIs numerically, append --signal-roi X,Y,W,H --background-roi X,Y,W,H. Coordinates start at 1,1; width and height are pixel counts. Add 1 to Fiji ROI x and y coordinates when transferring them.

**5.** Review records_v10.csv, frames_v10.csv, roi_v10.png, sweep_v10.png and status_v10.txt in the timestamped analysis_v10 folder. Python also saves configuration_v10.json, analysis_v10.npz and sweep_v10.svg. Check the listed warnings and whether the optimum is resolved (Section 8), and confirm the setting with fine sweeps and repeats.

#### B.2 Check a new configuration

Before analysing a new acquisition format, compare one frame with a manual signal-minus-background calculation. Inspect the saved ROI image, clipping code and image mode. Retain that check with the dataset.

SHA-256: 1442da722f30cf03639e8fd84cdbb26b7def28e82c9122a19b4be6b93c4807b9
